# A frequency-dependent mobilization of heterogeneous pools of synaptic vesicles shapes presynaptic plasticity

**DOI:** 10.1101/146753

**Authors:** Frédéric Doussau, Hartmut Schmidt, Kevin Dorgans, Antoine M. Valera, Bernard Poulain, Philippe Isope

## Abstract

The segregation of the readily releasable pool of synaptic vesicles (RRP) in sub-pool which are differentially poised for exocytosis shapes short-term plasticity at depressing synapses. Here, we used in vitro recording and modeling of synaptic activity at the facilitating mice cerebellar granule cell to Purkinje cell synapse to demonstrate the existence of two sub-pools of vesicles in the RRP that can be differentially recruited upon fast changes in the stimulation frequency. We show that upon low-frequency stimulation, a population of fully-releasable vesicles is silenced, leading to full blockage of synaptic transmission. A second population of vesicles, reluctant to release by simple stimuli, is recruited in a millisecond time scale by high-frequency stimulation to support an ultrafast recovery of neurotransmitter release after low-frequency depression. The frequency-dependent mobilization or silencing of sub-pools of vesicles in granule cell terminals should play a major role in the filtering of sensorimotor information in the cerebellum.

## INTRODUCTION

In neural networks, transfer of information largely relies on the ability of presynaptic terminals to transduce information encoded by changes in action potentials (APs) firing rate into changes in the release of neurotransmitters. During trains of APs, the immediate tuning of synaptic efficacy is determined by the combination of multiple parameters including AP frequency, past firing activities, number of release-competent synaptic vesicles (SVs), also referred as the readily-releasable pool (RRP), probability of release (*p_r_*), or the number of release sites (N). To date, the deciphering of cellular mechanisms underlying synaptic efficacy is challenged by a non-unified view of the identity of SVs belonging to the RRP (Neher, 2015). Depending on studies, the RRP may correspond to a large population of docked SVs releasable by hyperosmotic stimulation or at the opposite is restricted to SVs releasable by a single AP (Moulder and Mennerick, 2005; Pan and Zucker, 2009; Neher, 2015). Besides, there is increasing evidence showing that the RRP is constituted with heterogeneous population of SVs differentially poised for exocytosis and sequentially recruited before the fusion step. In the calyx of Held, the RRP can be separated into a fast-releasing pool (FRP) and a slowly releasing one (SRP) (Sakaba and Neher, 2001; Sakaba, 2006; Schneggenburger et al., 2012) and during a train of APs, SRP are first converted to the FRP and then maturate to a superpriming step before acquiring a high release rate (Lee et al., 2012, 2013). In the cerebellar cortex, at GC-molecular layer interneuron (MLI), a segregation of the RRP in two pools mobilized in a two-step process has also been described while with faster kinetics for the transition between the two steps (Ishiyama et al., 2014; Miki et al., 2016). Finally, these studies draw a model in which synaptic plasticity during high-frequency trains is shaped by the kinetics of mobilization of sub-pools of SVs in the RRP (Pan and Zucker, 2009; Miki et al., 2016). However, the way synapses manage these pools/steps upon a broad bandwidth of stimulation frequency are still unknown.

Counter intuitively, the size of the RRP does not predetermine the orientation of presynaptic plasticity during high-frequency train since the calyx of Held with a very large RRP (700 to 5000 SVs; Borst and Soria van Hoeve, 2012) immediately depress while GC to Purkinje cell (PC) synapses facilitates during tens of APs (Kreitzer and Regehr, 2000; Valera et al., 2012; Schmidt et al., 2013), despite a RRP combining a small size (4-8 SVs, Xu-Friedman et al., 2001) with a relatively high *p_r_* (Schmidt et al., 2013; Sims and Hartell, 2005; Valera et al., 2012 - but see Atluri and Regehr, 1996). Using computer simulations associated with variance analysis of postsynaptic responses, we previously showed that this striking facilitation is the consequence of an increase in *N* occurring in milliseconds upon a Ca^2+^-dependent process (Valera et al., 2012; Brachtendorf et al., 2015). Here, we investigated the cellular mechanisms underlying presynaptic plasticity at GC-PC synapses, not only during burst of activity but also upon fast changes in the stimulation frequency. We did reveal the existence of two pools of mature SVs with distinct releasable properties: the fully-releasable pool which can be released by a single AP and the reluctant pool available for fusion in a millisecond time scale upon high-frequency stimuli. Also, we showed that fully-releasable SVs can be specifically and almost completely silenced by low-frequency stimulation. The ultrafast recruitment of reluctant SVs underlies the large facilitation of glutamate release at high frequencies and the rapid recovery of a full capacity of glutamate release following the almost complete inactivation of fully-releasable SVs. We finally describe a model demonstrating how two pools of SVs can account for the low-frequency depression and the ultrafast recovery of glutamate release at high frequencies. These results support the idea that the presynaptic machinery of GC boutons acts as a dynamic filter of GC activities, and explain how a wide range of sensory inputs are computed within the cerebellar cortex.

## MATERIALS AND METHODS

All experimental protocols are in accordance with European and French guidelines for animal experimentation and have been approved by the Bas-Rhin veterinary office, Strasbourg, France (authorization number A 67-311 to FD)

*Slice preparation.* Acute horizontal cerebellar slices were prepared from male C57Bl/6 mice aged 18-25 days. Mice were anesthetized by isoflurane inhalation and decapitated. The cerebellum was dissected out in ice-cold ACSF bubbled with carbogen (95% O_2_, 5% CO_2_) and containing 120 mM NaCl, 3 mM KCl, 26 mM NaHCO_3_, 1.25 mM NaH_2_PO_4_, 2.5 mM CaCl_2_, 2 mM MgCl_2_, 10 mM glucose and 0.05 mM minocyclin. Slices were then prepared (Microm HM650V) in a ice-cold solution containing 93 mM *N*-Methyl-D-Glucamine, 2.5 mM KCl, 0.5 mM CaCl_2_, 10 mM MgSO_4_, 1.2 mM NaH_2_PO_4_, 30 mM NaHCO_3_, 20 mM HEPES, 3 mM Na-Pyruvate, 2mM Thiourea, 5 mM Na-ascorbate, 25 mM D-Glucose and 1 mM Kynurenic acid (Zhao et al., 2011). Slices 300 μm thick were briefly soaked in a sucrose-based solution at 34° c bubbled with carbogen and containing 230 mM sucrose, 2.5 mM KCl, 26 mM NaHCO_3_, 1.25 mM NaH_2_PO_4_, 25 mM glucose, 0.8 mM CaCl_2_ and 8 mM MgCl_2_ before being maintained in bubbled ASCF medium (see above) at 34° C until their use for experiments.

*Electrophysiology.* After at least 1 h of recovery at 34° C, a slice was transferred to a recording chamber. In order to block inhibitory transmission, postsynaptic plasticities, GABA_B_ and endocannabinoid signaling, slices were continuously perfused with bubbled ACSF containing blockers of GABA_A_, GABA_B_, NMDA, CB1 and mGluR1 receptors. To do so, the following antagonists were added: 100 μM picrotoxin, 10 μπι CGP52432 (3-[[(3,4-Dichlorophenyl)-methyl]amino]propyl(diethoxymethyl)phosphinic acid), 100 μM D-AP5 (D-(-)-2-Amino-5-phosphonopentanoic acid) and 1 μM AM251 (1-(2,4-Dichlorophenyl)-5-(4-iodophenyl)-4-methyl-N-(piperidin-1-yl)-1H-pyrazole-3-carboxamide) and 2 μM JNJ16259685 ((3,4-Dihydro-2H-pyrano[2,3-b]quinolin-7-yl)-(cis-4-methoxycyclohexyl)-methanone). Recordings were made at 34° C in PCs located in the vermis. PCs were visualized using infrared contrast optics on an Olympus BX51WI upright microscope. Whole-cell patch-clamp recordings were obtained using a Multiclamp 700A amplifier (Molecular Devices). Pipette (2.5-3 MQ resistance) capacitance was cancelled and series resistance (R_s_) between 5 and 8 mΩ was compensated at 80%. R_s_ was monitored regularly during the experiment and the recording was stopped when R_s_ changed significantly (> 20%). PCs were held at -60 mV. The intracellular solution for voltage-clamp recording contained 140 mM CsCH_3_SO_3_, 10 mM Phosphocreatine, 10 mM HEPES, 5 mM QX314-Cl, 10 mM BAPTA, 4 mM Na-ATP and 0.3 mM Na-GTP. Parallel fibers were stimulated extracellularly using a monopolar glass electrode filled with ACSF, positioned at least 100 μπι away from the PC to ensure a clear separation between the stimulus artifact and EPSCs. Pulse train and low-frequency stimulation were generated using an Isostim A320 isolated constant current stimulator (World Precision Instruments) controlled by WinWCP freeware (John Dempster, Strathclyde Institute of Pharmacy and Biomedical Sciences, University of Strathclyde, UK). The synaptic currents evoked in PCs were low-pass filtered at 2 KHz and sampled at 100 KHz (National Instruments).

*Simulation of transmitter release and Ca^2+^ dynamics.* Models for Ca^2+^-dependent SVs fusion and replenishment (Millar et al., 2005; Wölfel et al., 2007; Sakaba, 2008) were transformed into the corresponding ordinary differential equations and numerically solved using Mathematica 10.0 (Wolfram Research).

Release rates were obtained by differentiation of the fused state. Paired-pulse ratios were calculated from the ratios of release probabilities obtained by the integration of release rates. Parameters for the release sensor part of the sequential two-pool model (**Fig. 7A,** V, corresponding to the RRP) were similar to those used by Sakaba (Sakaba, 2008) with forward rate k_on_ = 1*10^8^ M^-1^s^-1^, backward rate k_off_ = 3000 s^-1^, cooperativity b = 0.25, and release rate γ = 5000 s^-1^. Parameters for the replenishment part of this model (R_0_, R_1_ representing the reluctant pool) were similar to those given by Millar et al. (2005) for a phasic synapse and found by trial and error. The forward and backward rate constants of Ca^2+^-dependent priming and unpriming were k_prim_ = 8*10^8^ M^-1^s^-1^ and k_unprim_ = 120 s^-1^, respectively. Ca^2+^-independent filling and unfilling rates were k_fill_ = 200 s^-1^ and k_unfil_ = 150 s^-1^, respectively. Ninety percent of released SVs were recycled into the unprimed pool, supported by a slow Ca^2+^-independent filling rate of k_basal_ = 0.002 s^-1^, reflecting filling from a reserve pool. Recovery from Low-Frequency Depression (LFD, see results section) was simulated by restarting the model at high frequency with the size of the releasable pool set to the value at the end of the 2 Hz train, and the unprimed pool (R_0_) fully recovered. The one-pool model **(Fig. 7F**) and the parallel two-pool model (**Fig. 7G**) and their parameters were taken from Wölfel et al. (2007).

Releases triggering Ca^2+^ signals were simulated as repeated Gaussian curves spaced by the interstimulus intervals and adjusted to match the amplitude (22.5 μM) and half-width (5 μs) of the estimated action potential-mediated Ca^2+^ signal at the release sensor of PF synapses (Schmidt et al., 2013). In the sequential two-pool model (**Fig. 7A**) SVs replenishment was driven by residual Ca^2+^ with an amplitude of 520 nM per pulse, dropping exponentially with a time constant of 42 ms and summing linearly depending on the length of the interstimulus interval (Brenowitz and Regehr, 2007). For the two other models (**Fig. 7G,F**) SV replenishment occurred at a constant rate of 2 SV/ms (Wölfel et al., 2007). Resting Ca^2+^ was assumed to be 50 nM.

*Data and statistical analysis.* Data were acquired using WinWCP 4.2.x freeware (John Dempster, SIPBS, University of Strathclyde, UK). Analyses were performed using PClamp9 (Molecular Devices), Igor (6.22A) graphing and analysis environment (Wavemetrics, USA). Error bars in figures show SEMs. Student’s *t* test or paired t-test were performed when data were normally distributed; the Mann-Whitney Rank Sum Test (MWRST) or the signed rank test were used in all other cases. Statistical tests were performed using SigmaPlot 11 (Systat Software). The levels of significance are indicated as *ns* (not significant) when *p* >0.05, * when *p* ≤ 0.05, ** when *p* ≤ 0.01 and *** when *p* ≤ 0.001.

## RESULTS

### Low-Frequency depression at the granule cell-Purkinje cell synapse

In order to better understand the mechanisms underlying the large facilitation of glutamate release during repetitive activities at the GC-PC synapse, we first analyzed the behavior of synaptic transmission over a broad range of frequencies within transverse cerebellar slices prepared from young mice (P17 to P25). To focus our study on presynaptic mechanisms and rule out the possibility of postsynaptic contribution, we pharmacologically blocked the induction of postsynaptic long term plasticity (LTP and LTD), NMDA-dependent plasticity (Bidoret et al., 2009; Bouvier et al., 2016), and endocannabinoid signaling (Marcaggi and Attwell, 2005; Beierlein et al., 2007). We then stimulated parallel fibers (PF) with trains of stimuli (50 pulses) elicited at different frequencies (2, 50 and 100 Hz) and at near-physiological temperature (34° C) (**Fig. 1A**). As already reported in rodent cerebellar slices (Kreitzer and Regehr, 2000; Valera et al., 2012; Schmidt et al., 2013), a sustained facilitation of synaptic transmission was observed up to the 25^th^-30^th^ stimulus at frequencies above 50 Hz, indicating that the number of SVs released increased with stimulation frequency. The paired-pulse facilitation (PPF) at the first interval increased with frequency (PPF EPSC_2_/EPSC_1_: 171.15% ± 10.03 at 50 Hz, *n*=9, 215.96% ± 6.90 at 100 Hz, *n*=27, **Fig. 1A).** After 30 to 40 stimuli, a depression (EPSC_n_/EPSC_1_ <1) was observed.

**Figure 1.**
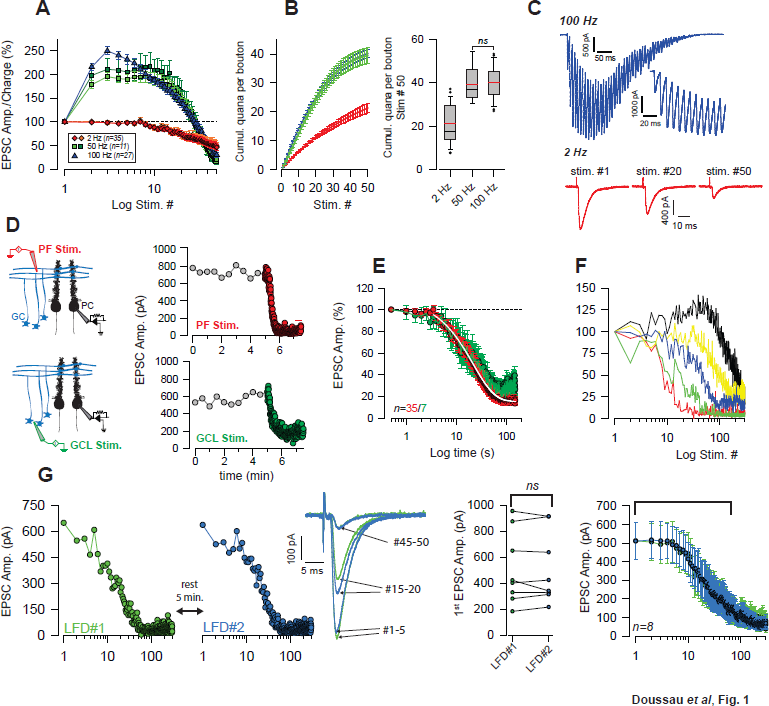
Low-frequency depression at GC-PC synapses. ***A,*** Averaged EPSC amplitude and charge versus stimulus number during train of stimuli at 2, 50 and 100 Hz. At 50 Hz EPSC amplitude and EPSC charge are represented with light green squares and dark green squares respectively. At 2 Hz, EPSC amplitudes and EPSC charge are represented with red diamonds and orange diamonds respectively. Due to the overlapping of EPSC, charge values were not measured at 100 Hz. Note the absence of facilitation at 2 Hz. ***B,*** Corresponding cumulative plots of the number of quanta release per bouton (left). The cumulative number of quanta released at 50 Hz and 100 Hz at the 50^th^ stimulus were statistically identical (*t*-test, *p*=0.82). Mean and median lanes are respectively represented in red and black (right) ***C,*** Examples of EPSCs during train at 100 Hz and 2 Hz. *Inset:* The first EPSCs observed during 100 Hz train application. The stimulus artifacts have been subtracted for the 100 Hz train. ***D,*** Time course of EPSC amplitude following stimulation of PF (upper panels) or GC somata (lower panels) at 0.033 Hz (gray points) and 2Hz (red or green points) ***E,*** Mean normalized EPSC amplitude during sustained 2 Hz stimulation of PF (red points) or GC somata (green points). Note the delay before the actual induction of depression. The depression was fitted by a monoexponential function (tau = 21.7 s, white lane). ***F,*** Selected time course of LFD recorded in 5 PCs showing differences in the onset and the plateau of depression ***G,*** Left graph: Time courses of two successive LFDs elicited in the same cell. The second LFD (LFD#2, blue points) was elicited after a resting period of 5 minutes after the first had ended (LFD#1, green points). Traces correspond to averaged EPSCs recorded during LFD#1 and LFD#2 (green and blue traces, respectively) at the indicated stimulus numbers. Middle graph: Values of EPSC amplitude recorded at the 1^st^ stimuli of LFD#1 (green points) and LFD#2 (blue points).The lack of statistical difference (*p*=0.94, paired *t*-test, *n*=8) in EPSC amplitudes between LFD#1 and LFD#2 reflects a full recovery from depression after the resting period that followed LFD#1. Right graph: Superimposition of mean EPSC amplitudes recorded during LFD#1 (green points) and LFD#2 (blue points). The similarity between LFD#1 and LFD#2 was tested using a paired t-test. The values were statistically different for the entire experiment (*p*<0.001, *n*=8) but the 60 firsts response were statistically identical (paired *t*-test, *p*=0.08, *n*=8).

At many synapses, asynchronous release of quanta can be triggered by repeated activities (Rudolph et al., 2011; Kaeser and Regehr, 2013). Since asynchronous release can be detected by changes in the charge/amplitude ratio, we checked whether the charge and peak amplitude of EPSC evolved in similar ways during 50 Hz and 2 Hz stimulation. The perfect superimposition of normalized charges and amplitudes at any stimulus number during 50 Hz and 2 Hz trains suggests that quanta released asynchronously did not participate in a detectable way during theses protocols (**Fig. 1A**). This enables us to estimate the number of quanta released during these trains. Values of quantal parameters found at unitary GC-PC synapses in mice (Schmidt et al., 2013) were used to estimate the number of quanta released per varicosity at GC-PC synapse during train a 2 Hz, 50 Hz and 100 Hz. The number of PFs stimulated in each experiment was estimated by dividing the mean value of EPSC amplitude at 0.033 Hz by the median EPSC amplitude obtained at unitary GC-PC synapse (5.3 pA, including release failure, Schmidt et al., 2013). The number of quanta released during each protocol was estimated by dividing the cumulative EPSC amplitude by the number of PF stimulated and by the mean value of the quantal content obtained at unitary GC-PC synapse (8 pA, Schmidt et al., 2013). Cumulative EPSC amplitude plots demonstrate that the total number of quanta released is proportional to stimulation frequency. However, although initial facilitation during the first two or three pulses is higher in 100 Hz train than in the 50 Hz train (Valera et al., 2012) no difference was observed in the maximal recruitment of releasable SVs between 50 Hz and 100 Hz (number of quanta released per bouton at 50 Hz = 39.47 ± 2.75, *n*=9 compared to 40.14 ± 7.50 at 100 Hz, *n*=27, *p*=0.82, *t*-test, **Fig. 1B**).

At 2 Hz, the synaptic responses failed to facilitate (**Fig. 1A,C**) and a sustained stimulation of PF during hundreds of stimuli led to a near full blockage of synaptic transmission (**Fig. 1D**). We named this rapid blockage of synaptic transmission “Low Frequency Depression” (LFD). Strikingly, the time course of EPSC amplitude in all the cells recorded (*n*=32) started to decrease mono-exponentially (**Fig. 1E**, mean time constant for depression at 2 Hz =15.9 s ± 1.2 s) after a median delay of 7 stimuli. It should be noted that a lack of LFD was observed for less than 5% of PC recorded in the vermis. These experiments were not taken in account for statistical analysis. Although it has already been shown that action potential is faithfully initiated and transmitted along parallel fibers at high frequencies and temperatures close to those in physiological conditions (35° C), (Kreitzer and Regehr, 2000; Isope and Barbour, 2002; Baginskas et al., 2009), repetitive extracellular stimulation of PF can induce decreases in fibers excitability via strong accumulations of K+ in the extracellular space (Kocsis et al., 1983). Hence, it could not be excluded that a decrease in the number of stimulated PF underpinned part of the blockage observed during sustained 2 Hz stimulation. To challenge this hypothesis, LFD was elicited by stimulating GC somata rather than PF (**Fig. 1D**). The lack of any significant change in the onset and the kinetics of depression after stimulating GC somata (**Fig. 1D-E**) suggested that change in PF excitability cannot underlie LFD. This hypothesis also draws on several lines of evidences presented later in figures 3, 4.

While the onset and the plateau of LFD were highly variable from one experiment to another (**Fig. 1F**), these parameters stay stable for a given PC as long as the recording of EPSCs could be maintained. In a series of 8 experiments, two LFD (LFD#1 and LFD#2) separated by a resting period of 5 to 10 minutes were successively elicited. As shown in figure 1 G, a full recovery of EPSC amplitude was achieved at the first stimulus of LFD#2 elicited after the resting period (paired *t*-test performed on EPSC#1 of LFD#1 and LFD#2, *p*=0.94, *n*=8). Strikingly, LFD#2 followed kinetics identical to LFD#1; the similarity between LFD#1 and LFD#2 were confirmed by comparing the mean values of EPSC amplitudes obtained at any stimulus number by a paired *t*-test. While LFD#1 and LFD#2 were statistically different when all stimuli were taken in account, we found no statistical differences for the first 60 stimuli (*p*=0.081, *n*=8) corresponding to the main time course of LFD (mean inhibition at stimulus #60: 66.8% ± 15.1%, *n*=8).

After pharmacologically blocking all known postsynaptic signaling events and therefore excluding a main postsynaptic contribution in LFD, these findings reveal for the first time that a sustained activity at low frequency can almost completely silences the release apparatus at GC terminals.

### Immediate recovery from LFD by high-frequency trains

Given the evidence that high-frequency trains of inputs can lead to the rapid refilling of empty sites or even the recruitment of new active release sites (Saviane and Silver, 2006; Hallermann et al., 2010; Valera et al., 2012; Chamberland et al., 2014), we tested whether a full capacity of release could be recovered after LFD by an increase in stimulation frequency. Alternatively, LFD may result from an activity-dependent blockage of presynaptic voltage-dependent calcium channels (Xu and Wu, 2005). In the latter case, accelerating the refilling of the RRP or increasing the number of release sites *N* should not restore glutamate release after LFD. LFD was induced by 300 stimuli at 2 Hz, and a train of 50 stimuli at 100 Hz was applied immediately after the last stimulus (**Fig. 2A**). After a full block of synaptic transmission, the recovery of glutamate release capacity occurred within 10 ms (**Fig. 2B,C**) and was sustained throughout a train of 50 stimuli (mean recovery within 10 ms: 70.7% ± 7.6). Strikingly, EPSC amplitudes in this 100 Hz recovery train rapidly reached values observed at the same stimulus number in a control train elicited before LFD induction (**Fig. 2C**), suggesting that the populations of SVs recruited in these 2 conditions are the same or share the same release properties (same size, same *p_r_*). Mean amplitudes of EPSCs recorded during train at 50 Hz and 100 Hz applied in control condition and after LFD induction were compared (**Fig. 2D**). Results showed that a full recovery from LFD was achieved within approximately 50 ms or 5 stimuli at 50 Hz (normalized EPSC amplitude at the 5^th^ stimulus: 203.67% ± 10.9 for trains applied in control condition compared to 165.00% ± 21.29 for trains applied after LFD, *p*=0.137, *t*-test, *n*=13) and 70 ms or 7 stimuli at 100 Hz (normalized EPSC amplitude at the 7^th^ stimulus: 207.24% ± 43.45 for trains applied in control condition compared to 191.12% ± 49.72 for trains applied after LFD, *p*=0.181, MWRST, *n*=27).

**Figure 2.**
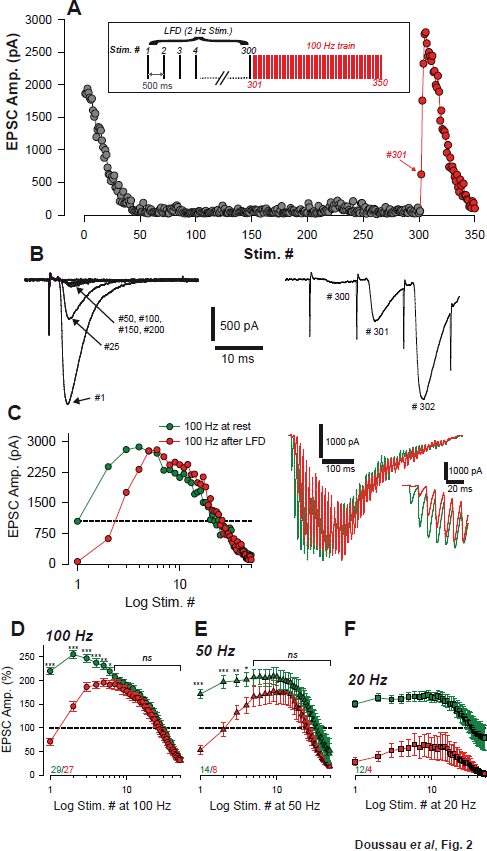
Ultrafast recovery from LFD by high-frequency trains. ***A,*** Typical experiment illustrating EPSC amplitude during LFD (gray points, stimulation at 2 Hz) and the fast recovery from LFD via high-frequency trains (red points, stimulation 100 Hz). *Inset:* protocol of stimulation. Stimulus #1 denotes the beginning of 2 Hz stimulation. ***B,*** Superimposed EPSC recorded during 2 Hz stimulation (left) and during the following 100 Hz train (right) at the indicated stimulus number. Note the ultrafast recovery from depression at 100 Hz (stimulus # 301). ***C,*** Left, Example of time course of EPSC amplitudes plotted against stimulus number during 100 Hz train applied in the control condition (green), or starting 10 ms after 300 stimuli at 2 Hz. EPSCs were recorded in the same PC. The dashed line corresponds to the baseline amplitude (mean value of EPSC at 0.033 Hz). Note the superposition of EPSC amplitudes after the 4^th^ stimuli at 100 Hz. Right, Corresponding traces recorded during these 100 Hz trains. *Inset:* The first EPSCs observed during train application. ***D,*** Mean values of normalized EPSC amplitude elicited by trains of stimulation at 100 Hz, 50 Hz and 20 Hz at baseline (0.033 Hz, green) or after LFD induction (2 Hz, red). EPSC amplitudes were not significantly different after the 7^th^ stimuli for 100 Hz trains (MWRST, *p*=0.181, *n*=27) and after the 5^th^ stimuli for 50 Hz trains (*t*-test, *p*=0.137, *n*=13). Numbers at the bottom of the graphs indicate the number of cells for each condition.

We then studied how the frequency of stimulation in the train influenced the ability of depressed synapses to recover a full capacity of release. Accordingly, 50 Hz and 20 Hz trains were applied after LFD and compared to 50 Hz and 20 Hz trains applied in control conditions. Mean amplitudes of EPSCs recorded during train at 50 Hz applied in control condition were not statistically different from those recorded after LFD induction (**Fig. 2E**), but at 20 Hz the recovery was limited (**Fig. 2F**). Two hypotheses can explain this ultrafast recovery of synaptic transmission following LFD: (1) a frequency-dependent ultrafast replenishment of the RRP, or (2) the recruitment of a reluctant pool of SVs that cannot be mobilized at low frequency as suggested in Valera et al. (2012).

### Recovery from LFD relies on the fast recruitment of reluctant SVs by a high-frequency train

In order to probe the existence of a reluctant pool supplying SVs to the fully-releasable pool at high frequency, we checked whether the recovery from LFD by a 100 Hz train could be affected by a previous exhaustion of this reluctant pool. As our previous results suggest that the reluctant pool can be recruited by short bursts at high frequency (paired-pulse/triplet stimulation at 50 or 100 Hz, Valera et al., 2012), we modified the paradigm of stimulation used to elicit LFD: LFD was induced by applying 100 triplets of stimulation instead of the 300 single pulses at 2 Hz (the total number of stimuli is the same in both paradigms). The triplet frequency (from 10 to 200 Hz) was changed during the experiment. This LFD was named LFD_triplet_. Since LFD protocols can be repeated several times after a recovery time (**Fig. 1F**), we systematically applied the LFD and LFD_triplet_ protocols to each PC recorded. Strikingly, although the number of stimuli was the same in both type of protocols (LFD classic or LFD_triplet_; **Fig. 3A, B**), recovery from depression was highly influenced by triplet frequency: the higher the frequency inside the triplets, the lower the level of recovery was seen to be (**Fig. 3B**). For example, with triplets at 200 Hz during LFD_triplet_, synaptic transmission hardly recovers (maximal recovery 38.9 % ± 7.19 *n*=6, compared to 196.7 % ± 8.7, after LFD, *n*=27, **Fig. 4C**), indicating that LFD_triplet_ depleted or disabled the population of SVs that support the recovery from LFD. If SVs 1) recruited only during LFD_triplet_ (at the second and third pulse of triplet stimulation) and 2) during recovery from LFD belong to the same pool (reluctant pool), then the number of quanta released during LFD and LFD_triplet_ should be the same. We performed a set of 18 experiments in which LFD and LFD_triplet_ (triplets at 100 Hz) successively applied in the same cells were both followed by the application of a 100 Hz recovery train (**Fig. 4A, B**). Cumulative EPSC amplitudes were determined, and the approximate number of quanta released per bouton for the whole protocol (LFD or LFD_triplet_ + recovery train) was estimated (**Fig. 4C**). The normalized cumulative EPSC amplitude was higher during LFD_triplet_ than during LFD (6440.9% ± 945% for LFD compared to 9353.5% ± 1199.9% for LFD_triplet_, *p*<0.001, paired *t* test, *n*=18, **Fig. 4C, D**). However, when the cumulative EPSC amplitude during LFD or LFD_triplet_ was summed with the cumulative EPSC amplitude of the recovery train, no further statistical differences were observed (11696.3% ± 1163.6 for LFD compared to 11379.0% ± 1349.6% for LFD_triplet_, *p*=0.61, paired *t* test, *n*=18, **Fig. 4C, D**). The number of quanta released during LFD was statistically lower than during LFD_triplet_ (42.7 ± 6.2 quanta per varicosity for LFD compared to 60.8 ± 8.0 quanta per varicosity for LFD_triplet_, *p*<0.01, paired *t*-test, *n*=18, **Fig. 4E**). However, when the number of quanta released during LFD or LFD_triplet_ was summed with the number of quanta released during the recovery train, no further statistical differences were observed (77.5 ± 7.7 quanta per varicosity for LFD + recovery compared to 74.8 ± 8.9 quanta per varicosity for LFD_triplet_ + recovery, *p*=0.51, paired *t*-test, *n*=18, **Fig. 4E**). Moreover, the number of quanta released during LFD and solely during recovery was statistically identical (42.7 ± 6.2 quanta per varicosity for LFD compared to 32.1 ± 2.3 quanta per varicosity for recovery, *p*=0.1, paired *t*-test, *n*=18, **Fig. 4E**).These experiments indicate that the RRP in GC boutons segregate in two distinct pools: the fully-releasable pool that can be released by a single action potential and a reluctant pool that can only be recruited during bursts of action potential reaching frequencies ≥ 50 Hz.

**Figure 3.**
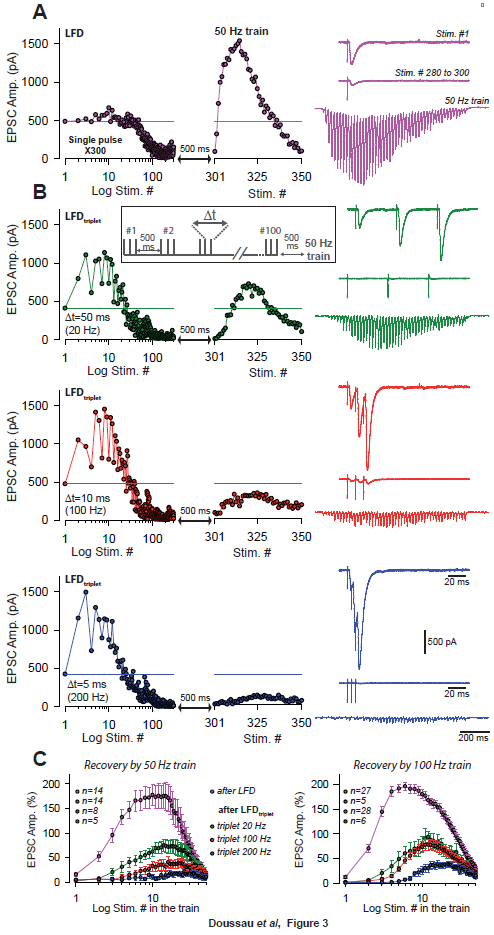
Short-term facilitation during triple-pulse stimulation at high frequency impedes recovery from LFD via high-frequency trains. ***A,*** LFD induced by a single pulse followed by a 50 Hz train ***B,*** LFDs induced by a triple pulse at different frequencies (LFD_triplet_, see inset for protocol), namely 20 Hz (green), 100 Hz (red) and 200 Hz (blue) followed by a 50 Hz train. For each condition, the upper traces correspond to EPSCs recorded at stimulus #1 (purple trace) or at triple pulse #1 (other traces) of the 2 Hz stimulation. The middle traces correspond to averaged EPSCs recorded during the LFD plateau (stimuli #280 to #300 for the purple trace and triple-pulses #80 to #100 for other traces). The bottom traces from which stimulus artifacts were subtracted correspond to EPSCs recorded during the 50 Hz train applied 500 ms after the 2 Hz stimulation ended. All data and traces were obtained in the same PC, and LFDs were elicited after a resting period of 5 minutes after the end of each protocol. ***C,*** Mean values of normalized EPSC amplitude plotted against stimulus number and recorded during 50 Hz train (left) and 100 Hz train applied 500 ms after LFD induction (300 single stimuli at 2 Hz or 100 triple pulses at 2 Hz). Color code is identical to that used in *A, B.*

**Figure 4.**
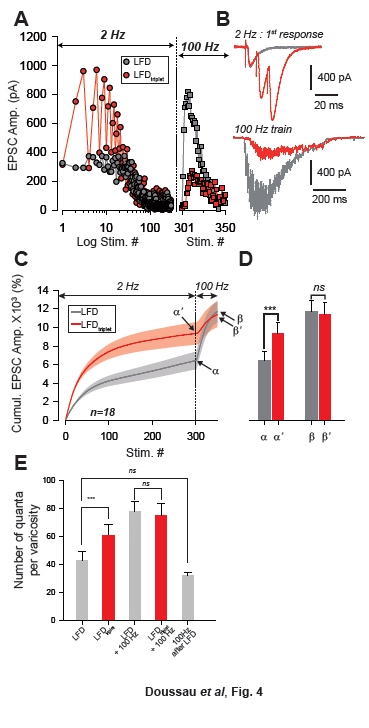
Recruitment of reluctant vesicles by high-frequency trains underpins recovery from LFD. ***A,*** Superimposition of the ESPC amplitude time courses elicited during LFD induction (gray circles) or LFD_triplet_ (triplet stimulation at 100 Hz, red circles) in the same PC, and during the recovery from LFD via application of a 100 Hz train (gray squares for a 100 Hz train applied 500 ms after LFD, and red squares for a 100 Hz train applied 500 ms after LFD_triplet_). ***B,*** The upper traces correspond to the superimposition of the first EPSCs recorded at stimulus #1 for LFD (gray trace) and LFD_triplet_ (red trace). The lower traces correspond to EPSCs recorded during the 100 Hz trains applied 500 ms after LFD (gray trace) or 500 ms after LFD_triplet_. ***C,*** Mean values of the cumulative EPSC amplitudes during LFD and LFD_triplet_ induction protocol followed by a recovery train at 100 Hz (n=18 cells). LFD and LFD_triplet_ were elicited successively in the same PCs. The dashed line indicates the beginning of the 100 Hz trains. α and a’ correspond to the cumulative value at the end of LFD protocols, β and β’ are values obtained following the recovery trains. ***D,*** Mean values of α, α’, β and β’ (same y axis as in C). ***E,*** Estimation of the number of quanta released per varicosity at GC-PC synapse during LFD, LFD_triplet_ and during the recovery via 100 Hz train (same set of experiments than in *D*). For the panels *D* and *E*, data were compared by using paired *t*-test.

While it may seem improbable, it may be argued that LFD and recovery by 100 Hz train could result from fast changes in PF excitability (Kocsis et al., 1983). However, the fact that 100 Hz trains are not able to reactivate synaptic transmission after LFD_triplet_ is another argument as to why this hypothesis cannot be retained.

### A two-pool model accounts for basic experimental findings

Our data suggest a segregation of SVs into different pools at GC terminals, which is in line with several previous studies (Valera et al., 2012; Ishiyama et al., 2014; Brachtendorf et al., 2015; Miki et al., 2016). They further suggest that SVs can be recruited from a reluctant pool into an fully-releasable pool on a millisecond timescale. Different previously published models could be suitable to account for these findings (**Fig. 5**), including a sequential two-pool model with Ca^2+^-dependent recruitment (Millar et al., 2005; Sakaba, 2008), a single-pool model with Ca^2+^-independent replenishment and a parallel two-pool model with intrinsically different SVs in which both pools are restored independently of Ca^2+^ (Wölfel et al., 2007; Schneggenburger et al., 2012). An important demand on such model is that it needs to account for our experimental results of high-frequency facilitation and LFD in a single model.

**Figure 5.**
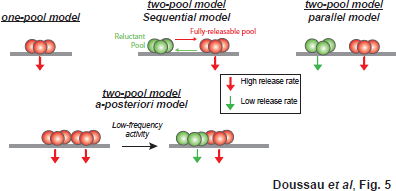
Schematics of the various models accounting for exocytosis at GC-PC synapses. Based on hypothesis proposed in other synapses, exocytosis could be achieved through an intrinsically homogeneous pool of release-ready SVs (one-pool model) or through a two-pool model with alternative stages of transition between the two pools. For the two-pool model, releasable SVs are separated in a fully-releasable pool (red SVs) or a reluctant one (green SVs).

First, we analyzed a sequential two-pool model of release and Ca^2+^-dependent recruitment (Millar et al., 2005; Sakaba, 2008). In this model, release is triggered via a Ca^2+^-driven, five-site sensor from a fraction of releasable SVs (**Fig. 6A**, see Methods). These SVs become replenished in two steps. The first step is a Ca^2+^-dependent priming step (R_0_→R_1_), followed by a Ca^2+^-independent filling step (R_1_→V), i.e. R_0_+R_1_ corresponds to the reluctant pool and V to the fully-releasable pool in our experiments.

**Figure 6.**
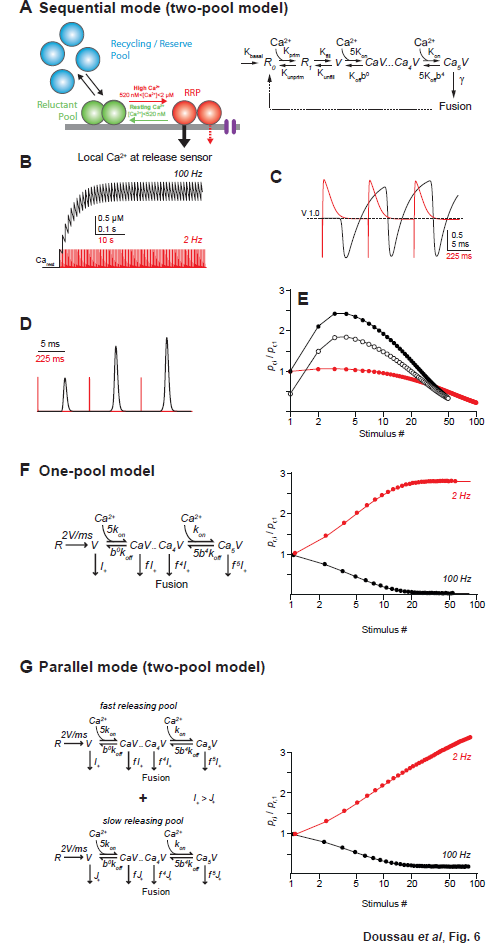
Models of facilitation and LFD. ***A,*** Scheme illustrating the sequential model of Ca^2+^ binding and release at the PF-PC synapse (left). Voltage-dependent calcium channels are represented in purple. In the corresponding mathematical model (right) the recycling / reserve pool is contained only implicitly by restoration of the reluctant pool (R_0_+R_1_) from fused SVs (dashed arrow) and via a basal refilling rate (*k_basai_*) Therefore, this model is referred to as sequential “two-pool” model. During high-frequency activation the residual Ca^2+^ increases, resulting in recruitment of SVs from the reluctant pool into the RRP, i.e. a temporal increase in the RRP that causes substantial facilitation (red arrow). The residual Ca^2+^ generates an additional moderate, short-lasting facilitation due to slow unbinding from the release sensor (dashed red arrow). During low frequency activation at 2 Hz, the residual Ca^2+^ fully drops back to resting level between stimuli and SVs recruited to the RRP return to the reluctant pool (green arrow). ***B,*** Simulated time courses and amplitudes of residual Ca^2+^ during high-(100 Hz, black) or low-frequency (2 Hz, red) activation starting from a resting Ca^2+^ (Ca_rest_) level of 50 nM. ***C,*** Fraction of Ca^2+^ unoccupied SVs in the RRP (V) of the release sensor during the initial three activations of a 100 Hz (black) or 2 Hz (red) activation train. Note that during the first three activations at low frequency the RRP relaxes to its initial (V=1, i.e. 100%) from a transient overfilling (V>1) prior to the next pulse while it continues to increase in size during high-frequency activation due to the build-up in residual Ca^2+^ and continuing recruitment of reluctant vesicles (cf. B). ***D,*** Transmitter release rates during three activations at high (black) or low frequencies (red), normalized to the first release process. ***E,*** Paired pulse ratios (PPRs) calculated as the ratio of release probabilities in the i-th (p_ni_) and the first pulse (p_r1_) during 100 Hz (filled black) or 2 Hz (red) activations plotted against the logarithmically scaled stimulus number. Open circles show PPRs duringh100 Hz activations started in a previously depressed model. ***F,*** (left) One-pool model of Ca^2+^ binding and release according to Wölfel et al (2007) consisting of the “allosteric” sensor model (Lou et al., 2005) supplemented with a reloading step of 2 SVs/ms. In contrast to our experimental findings, this model generates low-frequency facilitation and high-frequency depression (right). ***G,*** (left) as in ***F*** but for two parallel, non-interacting pools of SVs differing in their release rate constants thereby generating a “fast releasing pool of SV” (release rate I_+_ as in ***F***) and a “slowly releasing pool of SV” (release rate J_+_<I_+_ as in ***F***, Wölfel et al., 2007). Both models are restored via Ca^2+^ independent reloading steps of 2 SVs/ms. Note that similar to the model in ***F,*** this simulation generates a high-frequency depression and a low-frequency facilitation.

In the simulations, release was triggered by Gaussian shaped Ca^2+^ signals with amplitudes of ~22 (Schmidt et al., 2013). Ca^2+^-dependent recruitment was assumed to be driven by residual Ca^2+^ (**Fig. 6B**) with an initial amplitude of 520 nM and a decay constant of 42 ms (Brenowitz and Regehr, 2007). During high-frequency activation (100 Hz), this residual Ca^2+^ sums linearly (Brenowitz and Regehr, 2007), building up to a steady state amplitude of ~2 across 50 stimuli. This high concentration of residual Ca^2+^ drove rapid recruitment of SVs from the reluctant pool into the fully-releasable pool, leading to a transient increase in the size of this pool (**Fig. 6C;** Valera et al., 2012). Acting in concert with slow Ca^2+^ unbinding from the release sensor, which generates a moderate facilitation on the ms timescale (Bornschein et al., 2013), SV recruitment resulted in prolonged and facilitated release consistent with our experimental observations (**Fig. 6D, E**, compared with **Fig. 1**). During low-frequency activation (2 Hz), on the other hand, residual Ca^2+^ dropped back to its resting level between pulses and recruited SVs returned to the unprimed pool prior to the next stimulus. This resulted in the progressive depletion of fully-releasable SVs, leading to LFD. Renewed driving of the depressed model at high frequency reproduced a rapid recovery from the depression, and was followed by facilitation (**Fig. 6D** compared with **Fig. 2**), thus illustrating the recruitment of the reluctant pool.

Next, we simulated release using a single pool model consisting of the “allosteric” release sensor (Lou et al., 2005) supplemented with a replenishment step at a fixed, Ca^2+^-independent rate (one-pool model, Wölfel et al., 2007). Finally, we modeled release using two parallel pools of releasable SVs differing in their intrinsic release rate constants, thereby forming a pool of fast and a pool of slowly releasing vesicles (parallel two-pool model, Wölfel et al., 2007). Both pools were replenished independent of Ca^2+^ at a fixed rate as in the above single pool model. Notably, both the one-pool model and the parallel two-pool model predict a depression of release during high-frequency activation and a strong facilitation during sustained low-frequency stimulation (**Fig. 6F,G**). This is in stark contrast to our experimental results that showed the opposite behavior of GC synapses. Thus, although the latter two models successfully described several aspects of release at the depressing calyx of Held synapse that operates with a very large RRP (Sätzler et al., 2002), it appears that the small cerebellar cortical GC synapse utilizes different mechanisms to sustain release based on a small RRP and rapid, Ca^2+^ driven recruitment.

Taken together, these findings show that several aspects of our experimental findings, in particular high-frequency facilitation and LFD, can be reproduced by a simple sequential two-pool model with Ca^2+^-dependent recruitment but not, if recruitment is assumed to be Ca^2+^-independent. This does not exclude a contribution of more sophisticated mechanisms like activity dependent a posteriori modifications (Wölfel et al., 2007) or a separate facilitation sensor (Atluri and Regehr, 1996; Jackman et al., 2016) but hints towards the minimal requirements for sustained release from a small terminal operating with a small number of SVs in the RRP.

### Recovery from LFD depends on the size of both pools

The sequential model predicts that the kinetics of refilling of the fully-releasable pool after LFD depends on the size of in the reluctant pool. To test this hypothesis, we determined the recovery kinetics of the fully-releasable pool after LFD (**Fig. 7A**, red points) and LFD_triplet_ (triplet at 200 Hz; **Fig. 7A**, blue points) by applying a single stimulus at very low frequency (0.033 Hz) beginning 30 s after the end of the LFD protocol. As shown in Figure 7A, full recovery after LFD is described by a single exponential function (tau =153.8 s, R^2^=0.93), indicating a single-step process. However, recovery following LFD_triplet_ was biphasic: during an initial phase, release remained almost fully blocked. This initial phase was followed by a second phase that was again characterized by a single exponential time course (tau = 212.8 s, **Fig. 7A**). Averaged time courses suggest that the first phase could be considered as a delay, since a simple 90 s shift led to an identical monoexponential recovery in both LFD (mean tau =156.3 s, R^2^=0.98, *n*=8) and LFD_triplet_ (mean tau =149.3 s, R^2^=0.96, *n*=6) (**Fig. 7B,C**). This set of experiments suggests after depletion of both the reluctant and fully-releasable pools, the delay preceding the exponential phase of recovery from depression (~90 s) might reflect the time required to reconstitute the reluctant pool and then to refill the fully-releasable pool from the reluctant pool.

**Figure 7.**
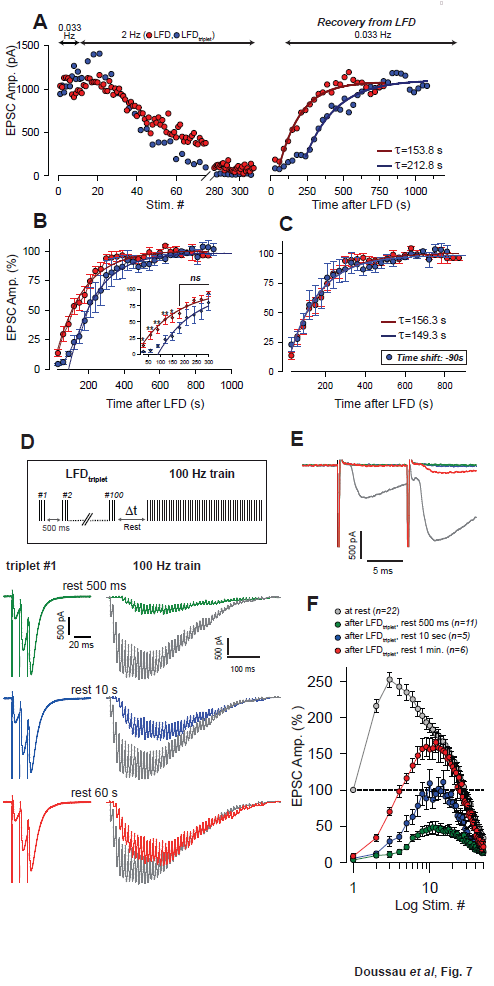
Kinetics of recovery from LFD ***A,*** Left, superimposition of the time course of LFD and LFD_triplet_ (triplet stimulation at 200 Hz) elicited successively in the same cell after at least 10 successive stimuli at 0.033 Hz. Right, time course for recovery from depression probed 30 s after the end of both LFD and LFD_triplet_ by a 0.033 Hz stimulation. For clarity, EPSC amplitudes were plotted against stimulus number for the left graph and against time for the right graph. The thick red and blue lines on the right graph represent the monoexponential fit of the recovery. Data could be fitted from the first EPSC recorded 30 s after the end of LFD, whereas the recovery after LFD_triplet_, could only be satisfactorily fitted after 7 stimuli at 0.033 Hz. ***B,*** Superimposition of the EPSC peak amplitude mean values plotted against time and recorded during a 0.033 Hz stimulation protocol applied 30 s after LFD (red points, *n*=8) or after LFD_triplet_ (triplet stimulation at 200 Hz, blue points, *n*=6). EPSC peak amplitudes were normalized to the mean value of EPSCs recorded during the plateau at 0.033 Hz. The thick red and blue lines on the right-hand graph represent the monoexponential fits. ***C,*** same values as in B, with the difference that the time axis was shifted by 90 sec for experimental values obtained during the recovery train applied after LFD_triplet_ (blue points). ***D,*** Recovery from LFD_triplet_ probed by 100 Hz train at the end of the LFD induction period, as indicated by the stimulation paradigm. Left, EPSC traces recorded during the 1^st^ triplet stimulation at 2 Hz. Right, EPSC traces recorded during 100 Hz trains applied 500 ms, 10 s or 1 min. after LFD induction ended. Traces are superimposed with EPSCs recorded during a 100 Hz train applied in control conditions (gray traces). ***E,*** Superimposition of the first responses to the trains recorded in *D. **F,*** Mean values of EPSC amplitudes recorded during 100 Hz trains applied 500 ms, 10 s or 1 min. after the 2 Hz stimulation period ended.

To check this hypothesis, the following paradigm was designed: SVs from both the reluctant and fully-releasable pools were released and inactivated by LFD_triplet_ (triplets at 100 Hz), then a 100 Hz test train was applied after a variable resting period (500 ms, 10 s or 1 min of rest after the end of LFD, **Fig. 7D**). The amplitudes of EPSCs recorded in the test trains were compared to those obtained from the application of a similar train in a control condition (before LFD). As shown in Figure 7D, the amplitude of EPSCs during the test trains increased proportionally to the length of the resting period. More interestingly, after 1 min of rest, the first responses in the test train were still fully blocked (**Fig. 7E**) whereas the amplitudes of the late responses (after stimuli # 11-12 in the test train) were similar to the corresponding responses in the control train (**Fig. 7F**). This demonstrates that after 1 minute of rest, no SVs from the fully-releasable pool were ready to be released, despite partial reconstitution of the reluctant pool and the replenishment of the fully-releasable pool by the reluctant pool.

### Reluctant SVS can be recruited by increasing Ca^2+^ entry

Finally, we investigated whether SVs in the reluctant pool are maintained in a low probability state. If so, the reluctant pool could be affected by manipulations known to affect *p_r_.* We therefore studied the properties of LFD and of its recovery by 100 Hz train after changes in *p_r_* or by affecting the spaciotemporal profile of [Ca^2+^] through the use of EGTA-AM (10 μM), a slow Ca^2+^ chelator that does not affect [Ca^2+^] in the nanodomain during single AP but that may impair the building-up of [Ca^2+^] during high-frequency train (Schmidt et al., 2013). Based on our work, increasing [Ca^2+^]_e_ from 2.5 mM (control condition) to 4 mM leads *p_r_* to increase from 0.25 to 0.67 for SVs in the fuly-releasable pool (Valera et al., 2012). In the presence of 4 mM [Ca^2+^]_e_, LFD time course was barely affected. Neither the delay (mean number of stimulus: 14.5 ± 2.8, *n*=30 at [Ca^2+^]_e_ = 2.5 mM compared to 17.6 ± 8.0, *n*=5, at [Ca^2+^]_e_, *p*=0.6, MWRST), nor the plateau of LFD (mean percentage of initial responses: 14.2% ± 2.0%, *n*=30 at [Ca^2+^]_e_ = 2.5 mM compared to 9.3% ± 1.4%, *n*=5, at [Ca^2+^]_e_=4 mM, *p*=0.69, MWRST) or the decay time constant (tau: 14.3 s ± 1.3 s, *n*=30 at [Ca^2+^]_e_ = 2.5 mM compared to 18.9 s ± 6.8 s, *n*=5, at [Ca^2+^]_e_ = 4 mM, *p*=0.87, MWRST) were statistically different in low and high Ca^2+^ conditions (**Fig. 8A, B**). At the opposite, recovery from LFD was strongly reduced (maximal recovery 85.5 % ± 11.6 *n*=5, **Fig. 8B)** high Ca^2+^ condition. This indicates that part of the reluctant pool was actually recruited during LFD upon increase in *p_r_.* Finally, we tested whether recruitment of the reluctant pool could be affected by impairing the spaciotemporal profile of [Ca^2+^]. Accordingly, LFD and the recovery form LFD by 100 Hz train were probed before and after application of 10 EGTA-AM. As previously showed, the application of 10 EGTA-AM did not affect the basal release of glutamate (**Fig. 8C**). Also, while LFD time course was insensitive to EGTA-AM (*p*=0,61, paired *t*-test performed on 3 successive EPSCs, *n*=9), the recovery from LFD was slightly slowed down and reduced during the first responses of the 100 Hz train; a statistical differences could be detected only at stimulus #4 and #5 of the train (*p*=0.031 for both stimuli, signed rank test, n=6) (**Fig. 8D**).

**Figure 8.**
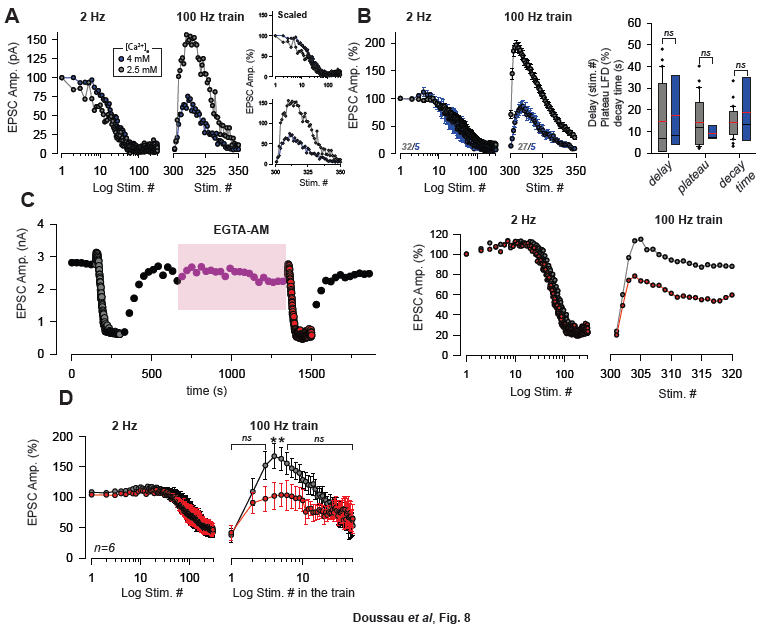
The reluctant pool can be recruited by increasing *p_r_. **A,*** Superimposition of the ESPC amplitude time courses, plotted against the stimulus number obtained in the same PC during LFD (left panels) and during the recovery from LFD (right panels) by a 100 Hz train before and after an increase in [Ca^2+^]_e_ from 2.5 mM to 4 mM. The inset shows the same experiment with normalized EPSC amplitude. ***B,*** Mean value of EPSC amplitude during LFD (left panel) and during the recovery via a 100 Hz train (middle panel) in presence of 2.5 mM (same set of data than in Fig. 1B, 3D) and 4 mM [Ca^2+^]_e_. Numbers at the bottom of the graphs indicate the number of cells for each condition. Right panel, the values of the delay (MWRST, *p*=0.60), plateau of LFD (MWRST, *p*=0.69) and decay time (MWRST, *p*=0.87) were statistically identical in low (2.5 mM, gray bars) and high Ca^2+^ (4 mM, blue bars) conditions. Black and red bars correspond to median and mean values respectively ***C,*** Left panel, typical experiment showing the time course of EPSC amplitude before and after the application of 10 EGTA-AM. Black points correspond to EPSCs recorded during 0.033Hz stimulation, gray and red points during 2 Hz stimulations and purple points during the application of 10 EGTA-AM at 0.033 Hz. Right panel, superimposition of normalized ESPC amplitudes from the same experiment during LFD and during the recovery via a 100 Hz train. **D,** Left panel, values of EPSC amplitude before (gray points) and after (red points) application of EGTA-AM are statistically identical (paired *t*-test, *p*=0.61, *n*=9) at any stimulus number during LFD. At the opposite, early responses were slightly reduced after application of EGTA-AM during recovery by 100 Hz train (*p*=0.031 at stimulus #5 and stimulus #6, signed rank test, *n*=6) (right panel).

## DISCUSSION

The present work demonstrates that the release of glutamate and presynaptic plasticity at the GC-PC synapse are shaped by the existence of 2 pools of releasable SVs, namely fully-releasable and reluctant SVs, bearing different properties for release. Whether both pools belongs or not to the RRP is still a matter of debate (Pan and Zucker, 2009; Neher, 2015) but if the RRP is restrained to SVs that can fuse with a single AP, as proposed recently (Miki et al., 2016), then it would correspond only to fully-primed SVs.

### Mechanism of recruitment of reluctant SVs

Our results showed that reluctant SVs that cannot be recruited during single stimuli at low-frequencies supports the release of glutamate during high-frequency train, even after a full depletion of fully-releasable SVs (**Fig. 2-4**). At GC-PC synapses, the localization of fusion events are restricted to distances that do not exceeds tens of nanometers from presynaptic Ca^2+^ entry even during short burst at high frequencies (Schmidt et al., 2013). The high speed of conversion (<10 ms) of reluctant SVs into fully releasable SVs suggests that reluctant SVs are docked in the nanodomain as a physical repositioning of reluctant SVs near the Ca^2+^-channels (positional priming) is probably too slow to occur with such fast kinetics.

The molecular mechanisms associated with the reluctant state and involved in the fast recruitment of reluctant SVs at GC-PC synapses are unknown, and are beyond the scope of this study. Our experiments suggest that fully-releasable SVs and reluctant SVs differ by their Ca^2+^ sensitivity and that 2 Ca^2+^ sensors differentially shape glutamate release during low- and fast-frequency stimulation (**Fig. 8**). In central synapses, the Ca^2+^ sensors synaptotagmin 1, 2 and 9 are involved in fast synchronous release triggered by a single AP (Xu et al., 2007) whereas synaptotagmin 7 which have slower kinetics for binding Ca^2+^ is involved in delayed asynchronous release (Wen et al., 2010; Luo et al., 2015). The recent finding that a lack of synaptotagmin 7 specifically impairs PPF without affecting *p_r_* (Jackman et al., 2016) makes this isoform as a good candidate for being involved in reluctant mechanisms. In another hand, Munc13s, actin and myosin II appears as good candidates for the fast recruitment of reluctant SVs. At the calyx of Held, actin is involved in the recruitment of reluctant SVs after the depletion of the FRP (Lee et al., 2012) and at GC-MLI synapses, actin and myosin II regulate the fast transfer of SVs from a “transition pool” to a “docked pool” (Miki et al., 2016). In the same synapses, the presence of Munc13s is required in a recently described “superpriming” step that would correspond to the final maturation stage of fully-competent SVs (Lee et al., 2013; Lipstein et al., 2013; Ishiyama et al., 2014). If we cannot exclude that identical mechanisms underpin the transition between both pools at the GC-PC synapse, one should keep in mind that this latter stands out by its large and sustained facilitation during high-frequency trains while, at the opposite, depression dominate upon high frequency stimulation at the calyx of Held and at some GC-MLI synapses (Schneggenburger et al., 1999; Bao et al., 2010). This suggests that both organization of SVs at the active zone and molecular mechanisms involved in the late stages of exocytosis are different or optimized at GC-PC synapses.

Both experimental data (**Fig. 7)** and *in silico* simulation (**Fig. 6)** suggest that reluctant SVs are recruited sequentially during high-frequency trains. Our model proposes that SVs can reversibly shift from the fully-releasable status to the reluctant one but do not explain why disequilibrium toward the reluctant state leading to LFD occurs at 2 Hz and not at slower frequencies. Hence, while some experiments clearly indicate that both pools are recruited sequentially (**Fig. 7**), one cannot exclude additional “a posteriori” mechanisms impeding the functioning of active release sites in a range of low frequencies (Wölfel et al., 2007; Schneggenburger et al., 2012). After SV fusion, release sites have to be purged of SV membrane and the time course of this clearance by endocytosis may act as a limiting factor for exocytosis during repetitive activities (Hosoi et al., 2009; Hua et al., 2013). In another hand, as proposed in invertebrates synapses (Silverman-Gavrila et al., 2005; Doussau et al., 2010), LFD may arise from an imbalance in the activation of presynaptic kinases and phosphatases and as such the kinase/phosphatase balance would act as a frequency sensor regulating the equilibrium between the two pools.

Our estimation indicates that the number of SVs that can be released during high-frequency trains after a full depletion of fully-releasable SVs corresponds to the size of the reluctant pool (**Fig. 5**). Since the number of SVs released per bouton (30-40 SVs, **Fig. 1, 5)** during 50/100 Hz trains, LFD, LFD_triplet_ or during recovery trains far exceeds the number of docked SVs counted in one varicosity at GC-PC synapses (4-8 SVs, Xu-Friedman et al., 2001), we postulate that the two releasable pools are refilled by other pools (recycling pool, reserve pool) during LFD, LFD_triplet_ and during the late phase of high-frequency trains. Previous works performed in other cerebellar synapses suggest that proteins of the cytomatrix (bassoon, actin, myosin II) (Hallermann et al., 2010; Hallermann and Silver, 2013; Miki et al., 2016) in concert with a Ca^2+^-driven acceleration of replenishment kinetics trains (Saviane and Silver, 2006) participate to the refilling of empty release sites during the last phase of high-frequency train.

### Segregation of releasable SVs in 2 pools shapes short-term plasticity

Neuronal networks in the cerebellar cortex have to process sensory information coded at ultra-high frequencies (up to 1 kilohertz, van Kan et al., 1993; Arenz et al., 2008, 2009). Most of this information is conveyed to the granular layer, the input stage of the cerebellar cortex, via the mossy fiber (MF) pathway. Strikingly, MF-GC synapses can sustain high frequency trains of input by using a specific arrangement of the presynaptic machinery to achieve ultra-fast reloading of SVs (Saviane and Silver, 2006; Rancz et al., 2007; Hallermann et al., 2010; Ritzau-Jost et al., 2014). However, debate continues over how these high frequencies inputs are integrated at the GC-PC synapses, the output stage of the cerebellar cortex and the major site for information storage in the cerebellum (Thach et al., 1992; Ito, 2006; D’Angelo and De Zeeuw, 2009). We already showed that none of the classical mechanisms for facilitation (including the buffer saturation model and residual Ca^2+^, Pan and Zucker, 2009; Regehr, 2012) that predict a pure increase in *p_r_* during PPF can satisfactorily account for the high PPF at GC-PC synapses and for its unusual ability to sustain glutamate release during tens of APs at high-frequency trains. During a train of APs at high frequency, the local [Ca^2+^] at release sites increases with AP number in the train (Schmidt et al., 2013; Miki et al., 2016) leading to an immediate increase in *N* that underlies high values of PPF (Valera et al., 2012; Brachtendorf et al., 2015). Here we showed that this increase in *N* arises from the fast recruitment of reluctant SVs into the fully-releasable pool. A fast and sequential recruitment of a “replacement pool” (analogous to the reluctant pool) in a docked pool (analogous to the fully-releasable pool) accounting for PPF have been recently described at GC-MLI synapses (Miki et al., 2016) but the GC-PC synapse is unique in so far that the sequential recruitment of both pools is associated with the possibility to specifically inactivate the fully-releasable pool by low-frequency stimulation. The combination of both mechanisms gives the striking possibility to filter repetitive activities around 2 Hz and to drastically invert the orientation of presynaptic plasticity (full depression versus strong facilitation) in response to changes in the stimulation frequency (**Fig. 2**). Finally, during high-frequency inputs, GC-PC synapses are able to reset and standardized synaptic efficacy independently of the recent history of previous events (**Fig. 2**).

### Physiological implications of the filtering of GC activity

These results provide new hypotheses about information processing in the MF-GC-PC pathway. *In vivo* experiments have shown that in lobules IV and V, GCs responding to joint rotation receive spontaneous synaptic inputs from MF leading to spontaneous firing at 2-10 Hz in GCs. Upon joint movement, GCs discharge shifts instantaneously in burst mode with frequencies from ~50 Hz to 300 Hz (Jörntell and Ekerot, 2006). Based on our work, we propose a model in which GC-PC synapses could act as a filter of GC activity. In the absence of sensory input, GC inputs to PCs would be filtered by an active silencing of presynaptic terminals. However, during joint movement, the efficient transmission of high-frequency activity from MF to GC (Saviane and Silver, 2006; Rancz et al., 2007; Hallermann et al., 2010) and from GC to PC (Valera et al., 2012), combined with the recruitment of the reluctant pool by high frequency trains at GC-PC terminals, would allow a reliable transmission of sensory information to PC. In such a model, the filtering of low-frequency activities originating from numerous GCs would enhance the signal to noise ratio and help PC to recognize relevant inputs generated by sensory inputs from spurious GC activities. Future studies could investigate the response of inhibitory transmission, and more specifically feed-forward inhibition, to the wide range of frequency observed in the GC firing pattern.

## ACKNOWLEDGMENTS

This work was supported by the Centre National pour la Recherche Scientifique, the Université de Strasbourg, the Agence Nationale pour la Recherche Grant (ANR-2010-JCJC-1403-1 MicroCer) and by the Fondation pour la Recherche Médicale to PI (# DEQ20140329514). We thank the TIGER project funded by INTERREG IV Rhin Supérieur program and European Funds for Regional Development (FEDER, # A31). This work was also supported by a DFG grant to HS (SCHM1838). AMV and KD were funded by a fellowship from the Ministère de la Recherche. AMV was also funded by a fellowship from the Fondation pour la Recherche Médicale.

We thank Sophie Reibel-Foisset and the animal facility Chronobiotron (UMS 3415 CNRS and Strasbourg University) for technical assistance. We thank Joanna Lignot (Munro Language Services) for proofreading.

